# Graph-based pangenome analysis uncovers structural and functional impacts of allopolyploidization events

**DOI:** 10.1101/2025.08.26.672342

**Authors:** Victor Loegler, Anne Friedrich, Joseph Schacherer

## Abstract

The emergence of graph pangenomes has significantly advanced the study of structural variants (SVs) across diverse species. However, the impact of complex polyploidization events, particularly allopolyploidy, on SV landscapes remains poorly explored. *Brettanomyces bruxellensis*, a yeast species shaped by multiple allopolyploidization events, offers a compelling model due to its well-characterized genomic and phenotypic diversity. Here, we integrated subgenome-resolved assemblies with population-scale genomic and transcriptomic datasets to characterize SVs distribution and their functional impact at the species level. Leveraging a constructed reference graph pangenome, we identified sequences that are unique to each subgenome, and detected 212,177 SVs across 1,060 sequenced isolates. Allopolyploid genomes harbored significantly more SVs (320 on average) than diploid genomes (116 on average), reflecting the extensive impact of hybridization on the genome architecture. A subset of SVs originating from ancestral divergence between subgenomes (hybridization-associated SVs) formed distinct hotspots of structural diversity. Furthermore, functional association analysis revealed that 27.5% of SVs exert local regulatory effects on gene expression. Interestingly, hybridization-associated structural variants disproportionately affect gene expression, with a greater proportion of associated variants and a significantly larger effect size compared to other SVs. Overall, these findings underscore the role of allopolyploidization in driving both structural and functional genomic diversification. Our study also highlights the value of graph-based approaches in dissecting complex genome evolution.

## Introduction

Polyploidization, defined as the duplication of whole genomes, is a major evolutionary force across eukaryotes, contributing to speciation, adaptation, and genomic innovation (*1–3*). One specific form, allopolyploidy, originates from the hybridization of related species, thereby introducing immediate genetic diversity by uniting divergent subgenomes within a single nucleus (*4*). The novel allelic combinations and structural rearrangements introduced by allopolyploidization may drive phenotypic evolution and ecological adaptation (*5–7*).

The consequences of polyploidization have been widely explored through the analysis of small genomic variants or gene sequences (*8*, *9*). Its impact on larger structural variants (≥50 bp), however, remains overlooked, partly due to the technical challenges associated with SV detection in polyploid genomes (*10*). Yet, the established functional impact of SVs in diploids indicates a potential for SVs to play a primary role in the diversification of polyploid species (*11–13*).

Recent advancements in long-read sequencing, haplotype-resolved assemblies and graph-based pangenomes now offer powerful tools to identify SVs in polyploid genomes (*14–20*). These approaches facilitate the clear segregation of haplotypes (or subgenomes), which is essential to track the structural consequences of allopolyploidization.

The budding yeast *Brettanomyces bruxellensis*, which has undergone multiple allopolyploidization events throughout its evolutionary trajectory, offers a valuable model to study this complex evolution of genomes (*21*). The species has been observed across a range of human-related environments and is mainly recognized for its role in beer and kombucha brewing, along with its role in wine spoilage (*22–26*). Five genetic clusters have been described in the natural population, including two with diploid genomes (the primary subgenome) and three with allotriploid genomes that carry each distinct acquired subgenomes in addition to the primary one (*27*). Moreover, substantial genomic and transcriptomic datasets are available for this species (*27–29*).

Here, we leverage these resources to investigate the structural and functional consequences of allopolyploidization. Using a newly constructed graph pangenome based on subgenome-resolved assemblies, we conducted a comprehensive analysis of the structural variant (SV) landscape in this species. We identified structural differences between subgenomes, revealed their concentration in specific genomic hotspots, and demonstrated their substantial impact on gene expression.

## Results

### A reference graph pangenome captures subgenome diversity

In a previous study, 71 long read genome assemblies with resolved subgenomes were generated, capturing the diversity of both diploid (D1, D2) and allopolyploid (A1, A2, A3) genetic clusters (*28*) (Fig. 1A). This collection of genomes, which includes 40 allopolyploid genomes, has been used to construct a reference pangenome graph. The Minigraph-Cactus algorithm (*15*) was employed to build the graph, starting from the primary linear reference genome(*30*). The resulting graph exhibits a total length of 26,926,529 bp, distributed across 5,464,509 segments. To assess the completeness of the graph, a rarefaction curve of the pangenome has been modelled using nonparametric asymptotic estimators. The total estimated length was determined to be 37.0 Mb, of which 72.8% is captured by our graph (Fig. 1B). According to the rarefaction model, a total of 169 isolates would be required to reach 90% of the species pangenome length (Table S1).

**Figure 1.**
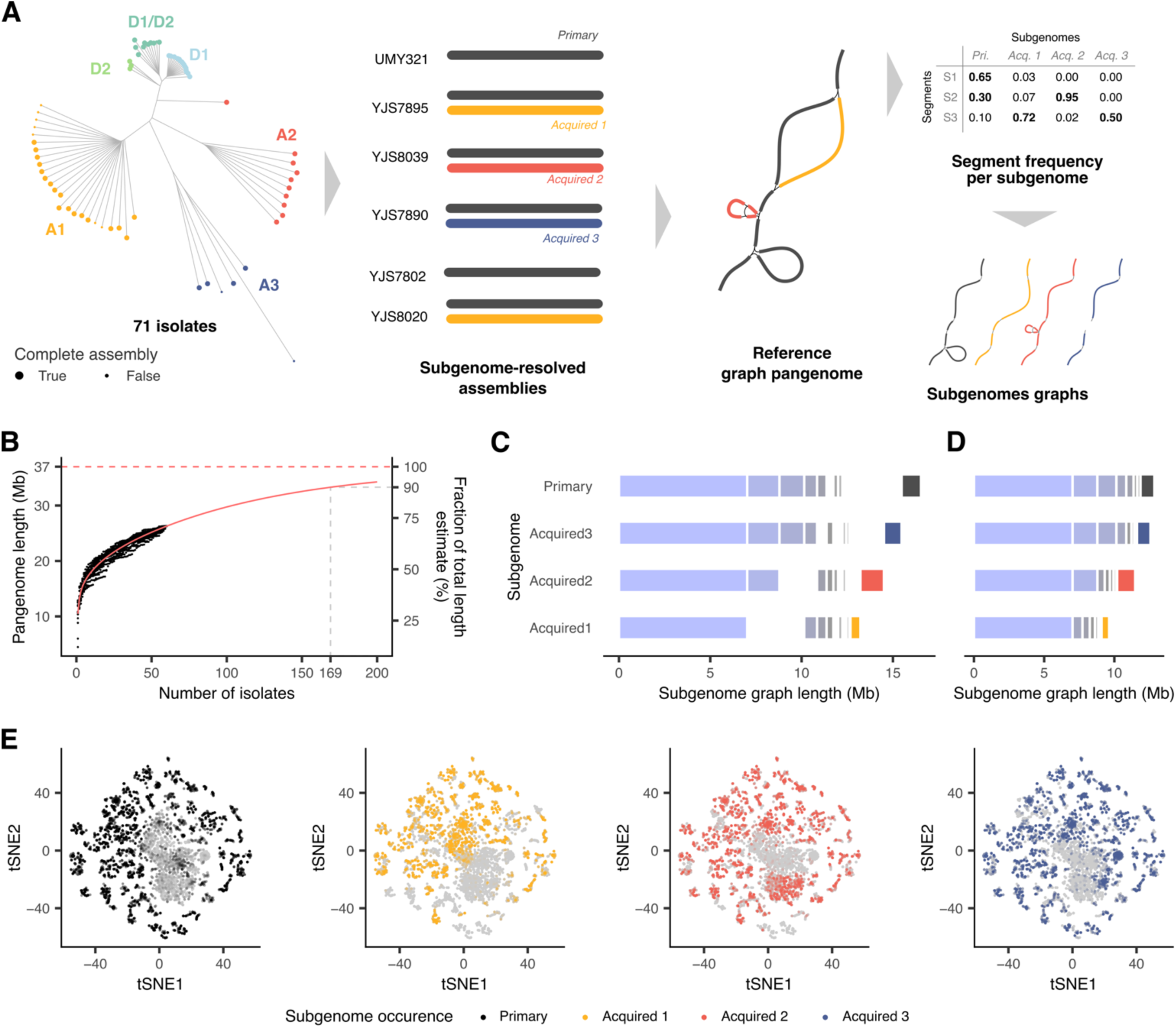
Reference graph pangenome. **A.** Neighbor-joining tree of the 71 isolates used for graph construction and method for subgenome graph identification. **B.** Rarefaction curve of the graph pangenome. The points represent observed rarefaction, the red curve correspond to the non-parametric rarefaction model and the red dashed line represents the estimated total pangenome length. **C.** Shared length of subgenome graphs. **D.** Total length of subgenome graphs. **E.** TSNE projection of segment occurrence across isolates. Segment occurrence within subgenome is color-coded.

To identify subgenomic patterns within the graph, we computed the frequency of occurrence of each segment within the primary and all three acquired subgenomes (Fig. 1A). Four subgenome-specific graphs were extracted by considering segments with 20% occurrence in the corresponding subgenome. The total length of subgenome-specific graphs ranges from 9.63, 11.49 and 12.58 Mb for acquired subgenomes 1, 2 and 3, up to 12.86 Mb for the primary subgenome (Fig. 1C). These results are consistent with the extent of acquired-to-primary genome conversion found in each allopolyploid genetic cluster, as converted regions are missing from the acquired subgenome contigs (*28*) (Fig. S1). A total of 7.02 Mb is shared between all subgenome graphs, and 0.55–1.3 Mb is private to each graph (Fig. 1C, Fig. S2, Table S2). A tSNE projection based on segment occurrence per isolate reveals clear segregation of subgenome-specific segments (Fig. 1D), revealing the complex genomic entanglement of the four subgenomes within the species.

### Primary subgenome drives long-term pangenome expansion

We quantified the length of non-reference sequences brought by each genome and subgenome. On average, genomes contain 853 kb of non-reference sequences. This number greatly varies between strains, with diploid isolates bringing 314 kb non-references sequences on average, while it goes up to an average of 1.3 Mb for allopolyploid isolates (Fig. 2A, Table S3). The acquired subgenomes bring a larger amount of non-reference sequence to the graph compared to the primary subgenomes (785 vs. 467 kb on average, two-sided rank-based Wilcoxon test, *P* = 1.65×10^-4^) (Fig. 2B, Table S3), which is consistent with their higher divergence from the reference genome.

**Figure 2.**
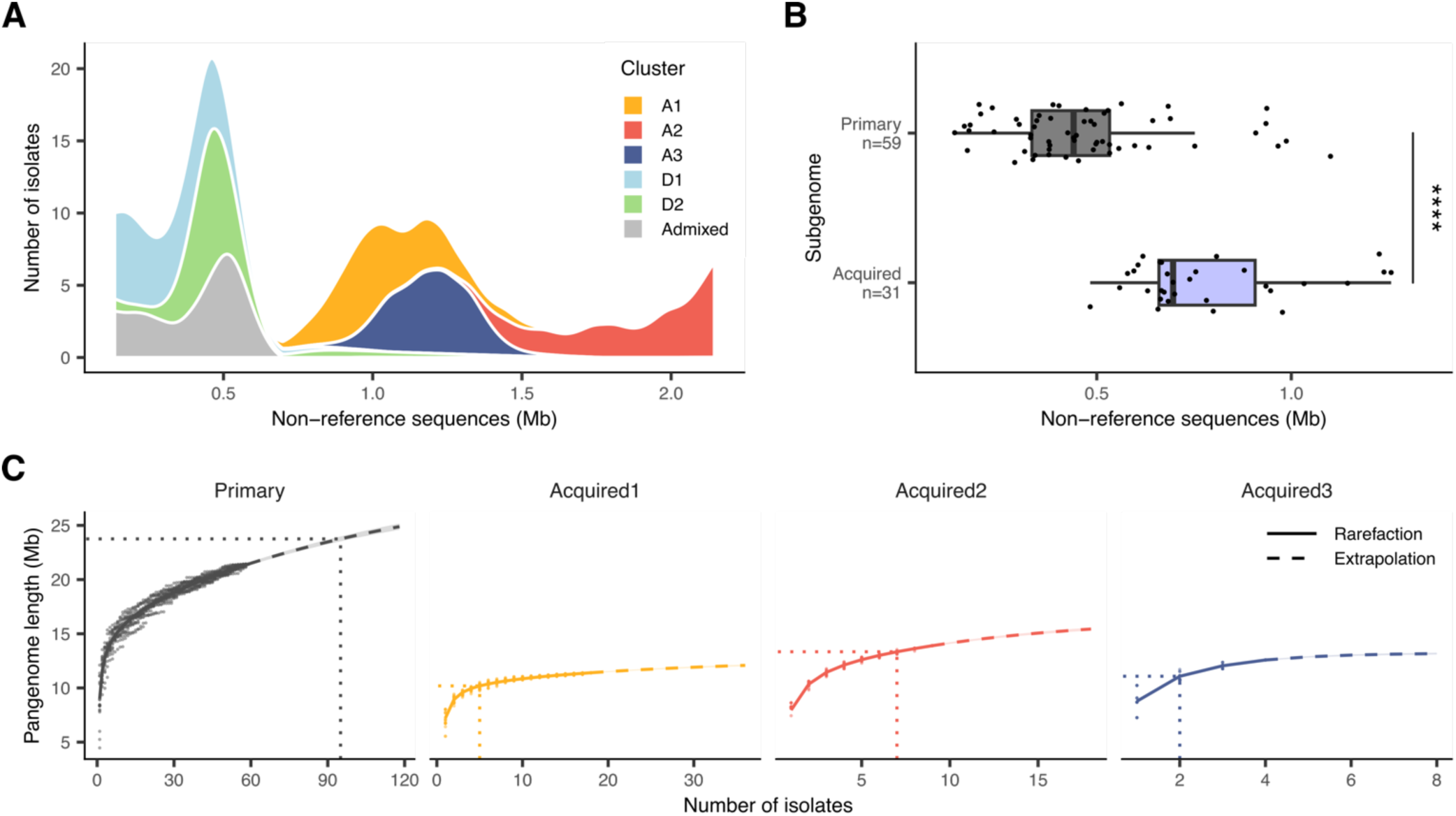
Pangenome expansion. **A.** Distribution of the amount of non-reference sequences found per genome colored by genetic cluster. **B.** Amount of non-reference sequences found per subgenome. The middle bar of the box plots corresponds to the median; the upper and lower bounds correspond to the third and first quartiles, respectively. The whiskers correspond to the upper and lower bounds 1.5 times the interquartile range (IQR). The *P* value was computed using a two-sided rank-based Wilcoxon test. **** = 5.7×10^-9^. **C.** Rarefaction curve per subgenome. The points represent observed rarefaction, the curves correspond to the non-parametric rarefaction model. The dotted lines indicate the pangenome length and number of isolates required to reach 80% of the pangenome saturation.

However, this early expansion rapidly reaches saturation in acquired subgenomes. Across all subgenomes in the population, 5, 7, and 2 isolates are required to reach 80% saturation of the A1, A2, and A3 pan-subgenomes, while 95 are required to reach this completion level for the primary pan-subgenome (Fig. 2C, Table S1). This marked contrast can be attributed, in part, to the presence of the primary subgenome across all genetic clusters, while the acquired subgenomes are cluster-specific. Nonetheless, even within allopolyploid clusters, more isolates are needed to reach saturation in the primary than in the acquired subgenome graphs. In the A1 cluster, 14 isolates allow to reach 80% of the primary genome estimated size, while this saturation is achieved with only 5 isolates for the acquired genome (Fig. S3A). Similar results are observed in the A2 cluster, where 10 and 8 isolates are required, respectively (Fig. S3B). These numbers reveal a lower diversity within the acquired genomes, hence a greater expansion of the primary genome as the number of isolates increases. This finding is coherent with previous observation of a lower level of diversity in the acquired subgenome in comparison to the primary one (*27*), and highlights the role of the primary subgenome in the long-term pangenome expansion.

### Species-wide structural variant mapping through a graph pangenome

The construction of a graph pangenome for *B. bruxellensis* has enabled the first analysis of the structural variant (SV) landscape of the species. A total of 212,177 redundant SVs were detected across a collection of 1,060 isolates (*27*) by mapping sequencing reads to the graph. This corresponds to 5,972 unique SV events, including 1,385 insertions, 939 deletions and 3,648 complex SVs (*Methods*). The length of SVs ranges from 50 bp to 18 kb, with an average length of 345 bp (Fig. S4). A substantial sequence redundancy is observed across SVs, with 655 insertions (47.3%) showing more than 90% sequence similarity with another (Fig. 3A). As such redundancy is often linked to the presence of transposable elements, we looked for the presence of miniature inverted-repeat transposable elements (MITEs) across all genomes. One MITE element has been previously reported in *B. bruxellensis* (*31*). Across the 71 genomes investigated, a total of 23 MITE families has been identified, with size ranging from 388 to 800 bp.

**Figure 3.**
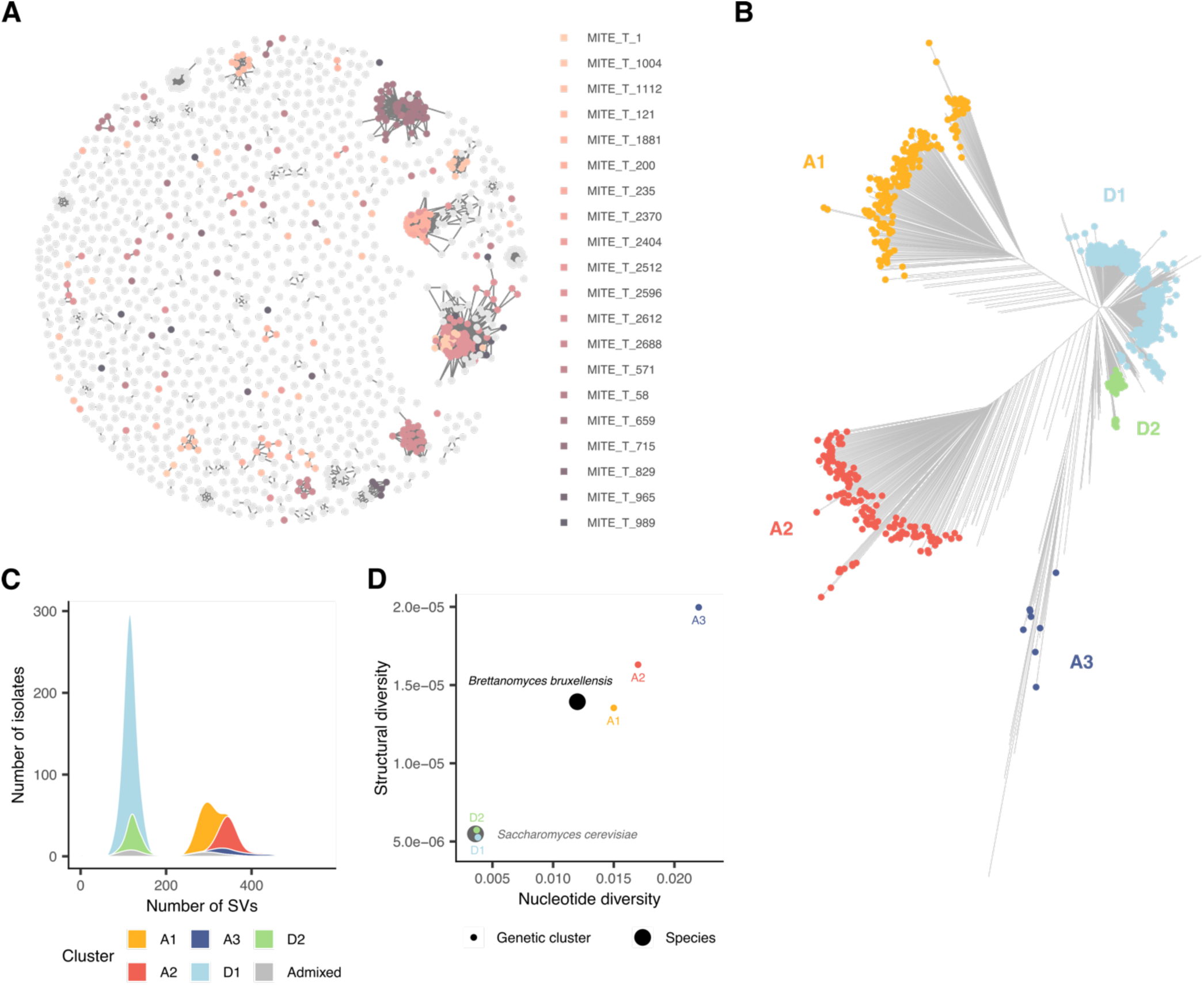
Structural variant landscape. **A.** Sequence similarity network of insertions. Each node represents one insertion, and edges indicate sequence identity ≥ 90% and ≥80% mutual coverage. MITE-related insertions are color-coded. **B.** Neighbor-joining tree based on 5,972 non-redundant SVs. The SNP-based genetic cluster are color-coded. **C.** Distribution of the number of SVs per isolate, colored by genetic cluster. **D.** Nucleotide and structural diversity of large collections of isolates for *Saccharomyces cerevisiae*, *Brettanomyces bruxellensis*, and genetic clusters of *B. bruxellensis*.

Subsequent comparison with SV sequences enabled the characterization of 1,092 MITE-related SVs, constituting 18.3% of the total number of SVs detected. The presence of MITE-related SVs explained nearly all the sequence redundancy observed within insertion sequences, suggesting that no other major transposable elements are related to SVs in the species (Fig. 3A).

### Allopolyploidy enhances structural variation in the population

The distribution of SVs in a population generally mirrors the pattern observed for smaller genomic variants (*32–34*). To verify this statement in our population, an SV-based neighbor-joining tree was constructed and further compared with the genetic cluster previously established on a SNP-based analysis (*27*). The obtained phylogeny is in complete accordance with the genetic clusters (Fig. 3B). Furthermore, as with small variants, the number of SVs found in each isolate deeply varies between non-allopolyploid and allopolyploid states. While the former exhibits an average number of 116 SVs, the latter has an average number of 320 SVs (Fig. 3C) (two-sided ranked-based Wilcoxon test, *P* < 2.2×10^-16^). In addition, the average proportion of heterozygous SVs is 0.70 for allopolyploid isolates in comparison to 0.46 for non-allopolyploid isolates (Fig. S5). This represents a significant difference (two-sided ranked-based Wilcoxon test, *P* < 2.2×10^-16^), as expected by the inclusion of an additional distant haplotype in allopolyploid genomes.

To compare the level of structural diversity with a non-allopolyploid yeast species, we took advantage of a reference graph pangenome constructed using 500 haplotypes of *Saccharomyces cerevisiae* (*35*) and used the same genotyping strategy within a collection of 3,034 natural isolates (*36*). The structural diversity, measured as the per-base average number of SVs between pairs of sequences (*Methods*), had a value of 1.39×10^-5^ in the *B. bruxellensis* population. The *S. cerevisiae* population, despite containing threefold more isolates, only exhibited a structural diversity of 5.49×10^-6^, which is 0.4-fold that found in *B. bruxellensis*. This mirrors the observed difference in nucleotide diversity:

1.2×10^-2^ for *B. bruxellensis* (*27*) and 3.6×10^-3^ for *S. cerevisiae* (*36*) (Fig. 3D). The structural diversity was further computed within each genetic cluster. The A1, A2, A3, D1 and D2 clusters exhibit a structural diversity of 13.5, 16.2, 19.5, 5.0 and 5.6×10^-6^, respectively (Fig. 3D). As for the nucleotide diversity (*27*), the structural diversity of allopolyploid cluster is higher than that of diploid clusters (two-sided ranked-based Wilcoxon test, *P* < 2.2×10^-16^). The level of structural diversity of non-allopolyploid clusters is of the same order of magnitude than the structural diversity of *S. cerevisiae* (Fig. 3D), indicating allopolyploidization as a main driver of structural diversity.

### Hotspots of structural variants arise from ancestral subgenome divergence

By adding novel divergent haplotypes into a population, hybridization participates to an increase in structural diversity. SVs previously fixed between the two parental species are now polymorphic and heterozygous in the resulting population. We took advantage of the subgenome-aware graph pangenome to segregate SVs resulting from mutation and propagation in the population, and those gained through hybridization, which correspond to ancestral structural rearrangements between subgenomes. The latter can be identified by their prevalence in one of the three acquired genomes (allele frequency – AF > 0.75) and low frequency in the primary genomes of the corresponding isolates (AF < 0.05). A subset of 31 allopolyploid isolates with qualitative primary and acquired genome assemblies was considered for this analysis. The distribution of allele frequency demonstrates a clear bias towards high frequencies for SVs located on the acquired genomes (Fig. 4A; two-sided rank-based Wilcoxon test, *P* < 2.2×10^-16^). Using the previously mentioned frequency thresholds, we identified 278 SVs (4.8%) introduced through hybridization (hereafter referred to as hybridization-associated SVs) and corresponding to ancestral SVs between subgenomes. Within hybridization-associated SVs, 114, 135, and 35 were observed as ancestral difference between the primary subgenome and the A1, A2, and A3 subgenomes, and only 5 SVs were brought by multiple acquired subgenomes. Despite the limited number of hybridization-associated SVs detected, their large frequency within the population, and their uneven distribution along the genome, result in considerable hotspots of structural diversity, associated with ancestral structural differences between subgenomes (Fig. 4B).

**Figure 4.**
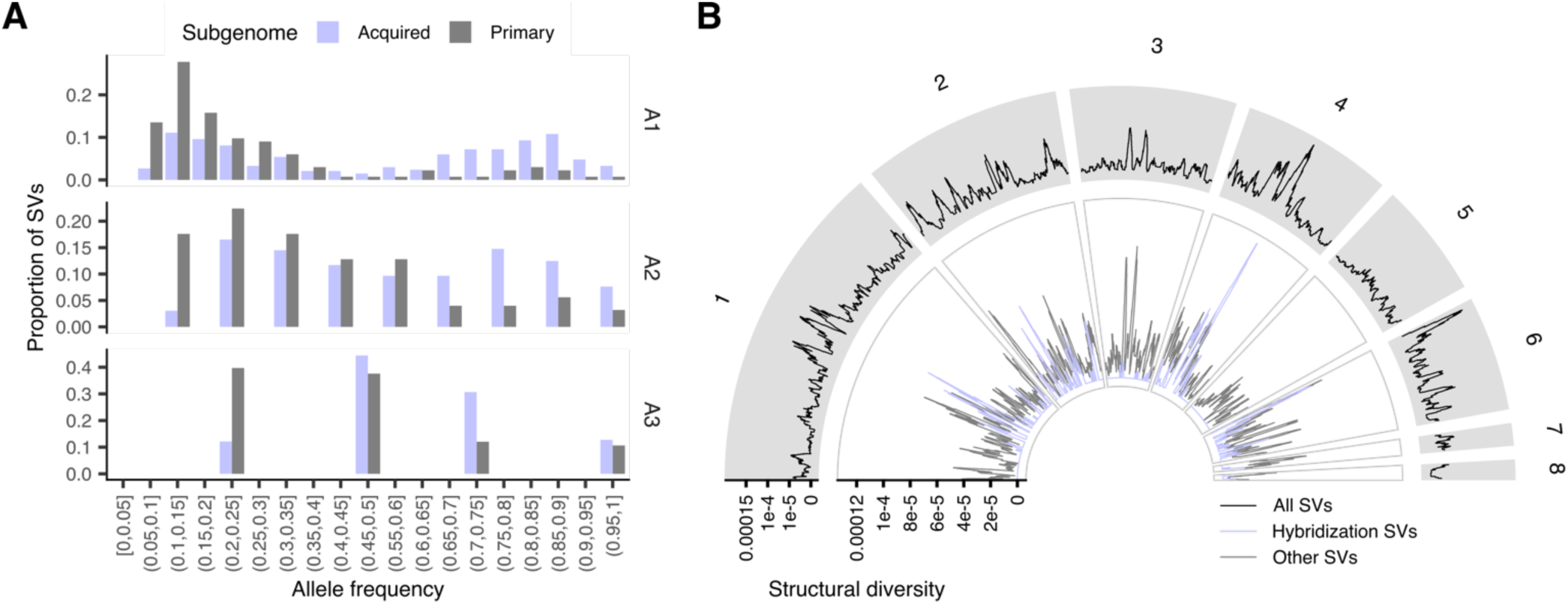
Distribution of hybridization SVs. **A.** Distribution of allele frequency per subgenome in the three allopolyploid clusters, using non-singleton SVs. **B.** Structural diversity along the genome using all SVs (outer ring), hybridization-associated SVs (inner ring, purple line) and other SVs (inner ring, gray line).

### Hybridization-associated structural variants disproportionately affect gene expression

Finally, we sought to investigate the local effect of SVs on gene expression by leveraging a transcriptomic dataset generated and available for diverse 87 isolates (*29*). A total of 32,590 SV-gene pairs separated by less than 50 kb were considered for association testing, involving 4,952 genes and 857 SVs. Using a linear mixed model to prevent the confounding effect of population structure, 436 SV-gene pairs showed significant association (Fig. 5A, Table S4). Out of the 857 SVs tested, 236 (27.5%) have a local effect on gene expression, affecting 298 genes (6.0% of all genes tested). Within associated SVs, there are 132 deletions (36.1% of all deletions tested), 53 insertions (26.5%) and 51 complex SVs (17.5%). Deletions are more frequently associated than insertions (two-sided Fisher’s exact test, odds ratio = 1.56, *P* = 0.024) and complex SVs (two-sided Fisher’s exact test, odds ratio = 2.65, *P* = 1.2×10^-7^). Despite a limited sample size (n=6), we observe a tendency for SVs overlapping genes to have a higher effect size on gene expression than SVs located upstream and downstream the gene of interest (Fig. 5B) (two-sided ranked-based Wilcoxon test, *P* = 0.021 and 0.022, respectively).

**Figure 5.**
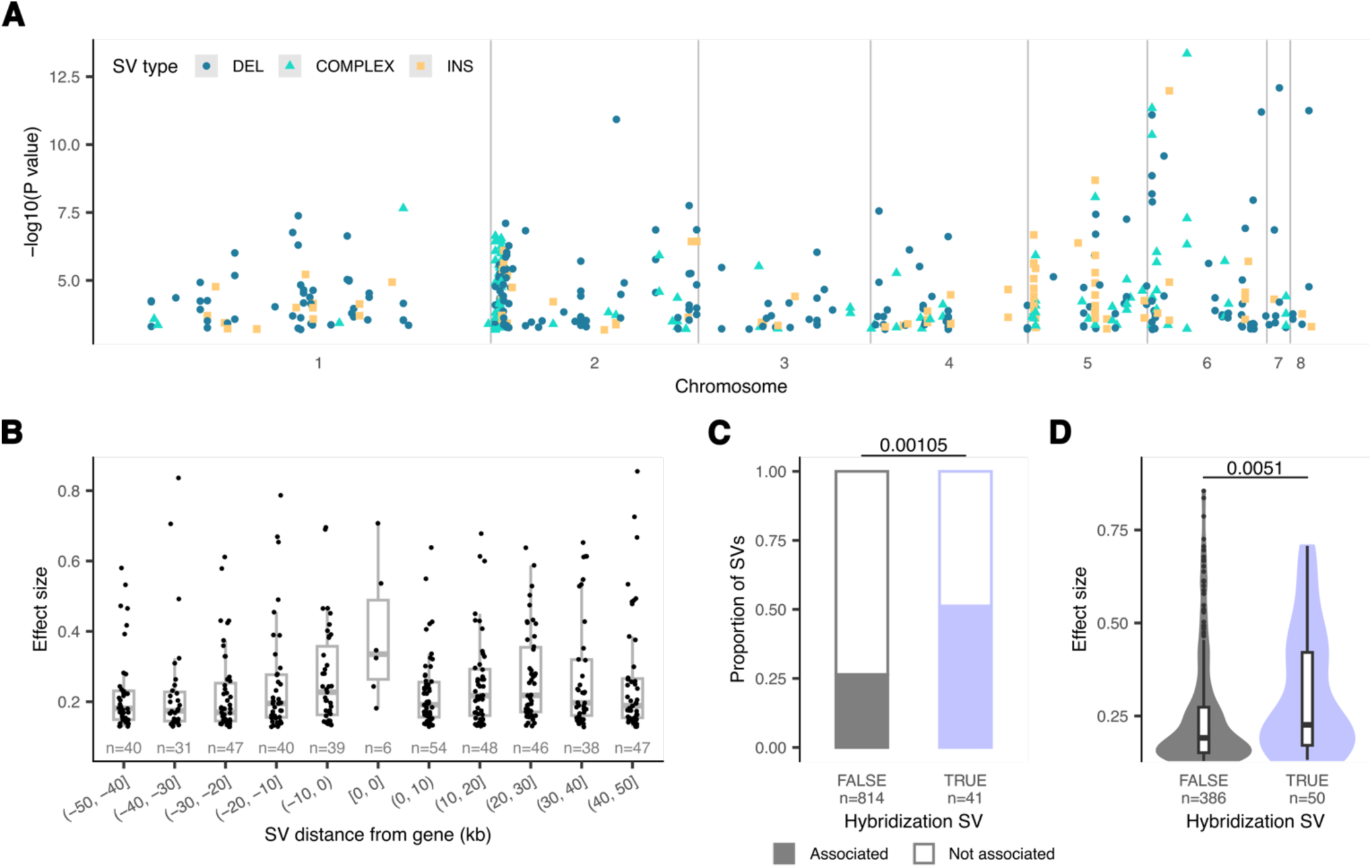
SV associations with gene expression variation. **A.** Associated SVs along the genome. The type of SV is color- and shape-coded. The x axis represents the -log10(*P* value) after FDR adjustment. **B.** Effect size depending on the distance of the SV in relation to the gene whose expression is affected. The middle bar of the box plots corresponds to the median; the upper and lower bounds correspond to the third and first quartiles, respectively. The whiskers correspond to the upper and lower bounds 1.5 times the interquartile range (IQR). **C.** Proportion of associated SVs among hybridization and other SVs tested for association. The *P* value was computed using a two-sided Fisher’s exact test. **D.** Effect size of hybridization SVs and other SVs. The middle bar of the box plots corresponds to the median; the upper and lower bounds correspond to the third and first quartiles, respectively. The whiskers correspond to the upper and lower bounds 1.5 times the interquartile range (IQR). The *P* value was computed using a two-sided rank-based Wilcoxon test.

To further explore the impact of allopolyploidization, hybridization-associated SVs were identified across all SVs tested for association using a competitive mapping strategy across subgenomes (*Methods*). A total of 40 were identified as hybridization-associated SVs, and 21 (51%) of these were associated with local differential gene expression. This represents a significant enrichment of hybridization-associated SVs (Fig. 5C; odds ratio = 2.92; two-sided Fisher’s exact test, *P* = 0.00105). In addition, hybridization-associated SVs exhibit a higher effect size than other associated SVs (0.31 vs. 0.24 on average; two-sided rank-based Wilcoxon test, *P* = 0.0051; Fig. 5D). Overall, this suggests that allopolyploidization have a substantial impact on gene expression diversification through the emergence of structural diversity.

## Discussion

Polyploidization events have profound consequences on species genomic diversity. In particular, polyploidy can increase structural diversity within populations in two ways. First, increasing the DNA content of cells amplifies the mutational input, as well as the masking of recessive deleterious variants. This is expected to result in a higher diversity in polyploids than in diploids. Consistently, recent studies demonstrated a higher SV load in tetraploid genomes than in diploid genomes in the plant genus *Cochlearia* and *Arabidopsis* (*37*, *38*). Second, in the context of allopolyploidization, the integration of a divergent genomic copy immediately increases structural variation. Structural differences fixed in the genomes of each parental species become suddenly part of the SV repertoire. Here, we demonstrated that these hybridization SVs strongly affect gene expression and may largely contribute to the phenotypic diversification of allopolyploid isolates (*27*).

Graph-based pangenomes played a pivotal role in enabling these findings. Although pangenomes are now routinely performed for diploid species, polyploid pangenomes remain scarce (*19*, *20*, *37*, *39*). Nevertheless, graph pangenomes built with appropriate phased genomes offer powerful tools to explore structural genome evolution in polyploids. For instance, the allopolyploid peanut pangenome revealed subgenome-biased structural diversity (*20*), and the potato pangenome revealed low haplotype diversity attributed to a recent population bottleneck (*19*), similar to that observed in *B. bruxellensis* acquired subgenome.

By generating a reference graph-based pangenome, this work also constitutes a valuable resource for further investigation of SVs in the species. Given the industrial significance of *B. bruxellensis* in beer and wine fermentation (*40–42*), our graph may support the characterization and selection of isolates for specific applications .

Despite these advances, two major limitations should be acknowledged. First, the subgenome phasing method relies on sequence divergence (*28*), which is highly efficient to segregate parental subgenomes but may miss inter-subgenome recombination events. More accurate phasing using long-range technologies, such as Hi-C, could resolve this issue and uncover additional structural complexity. Second, our current graph pangenome captures ∼75% of the estimated species diversity and could be improved by the addition of new diverse genomes.

Overall, this study highlights the structural and regulatory consequences of allopolyploidization, while demonstrating the power of graph pangenomes to elucidate genome evolution in complex systems.

## Methods

### Graph pangenome construction

Long-read, subgenome-resolved assemblies were retrieved for 71 isolates from a previous study (*28*). To minimize the number of false positive variants, the genomes were polished using paired-end Illumina sequencing data (*27*) with Hapo-G v1.3.2 (*43*). For each allopolyploid isolate, subgenome assemblies were concatenated into a single fasta file, polished, and contigs belonging to each subgenome were separated further. The graph pangenome reference was generated using Minigraph-Cactus v2.6.4 (*15*) with options *--reference UMY321 --vcf --gfa --gbz*, starting from the primary subgenome reference (*30*). Each was then added in a random order. For non-allopolyploid isolates, the primary genome was attributed to both haplotypes. For allopolyploid isolates, the acquired genome was attributed to the first haplotype, and the primary subgenome to the second.

### Graph pangenome processing

The graph pangenome was converted into a presence absence matrix with contigs as columns and segments as rows, using the GFA file from Minigraph-Cactus output. Briefly, a custom Python script was used to retrieve the presence of each segment in each contig from the walk lines in the GFA file. Since the subgenome and isolate corresponding to each contig can be traced back, this matrix allows one to check for the presence of segment per contig, subgenome, or isolate. Of the 71 isolates used for graph pangenome construction, a subset of 59 isolates with complete genome assemblies were retained (defined as having both an acquired and a primary subgenome longer than 8 Mb, which corresponds to the estimated size of the shortest acquired subgenome in the population). Subgenome-specific graphs were extracted by filtering out fragments that were present in less than 20% of the corresponding subgenome. For calculations requiring extensive computation time (*e.g.*, tSNE and rarefaction modeling), a random subset of segments was used, and the results were scaled to match the actual pangenome length.

### Rarefaction modelling

The R package iNext (v3.0.0) (*44*) with options *k = 400* and *size = seq*(*1, 300*) was used to model the rarefaction of the pangenome length with nonparametric asymptotic estimators. The modeling was performed using the presence-absence matrix of 10,000 nucleotides among isolates, and scaled back to the actual pangenome length. For subgenome rarefaction modeling, only contigs attributed to the corresponding subgenome were retained.

### Structural variant calling

The variant matrices were obtained in two ways. First, the Minigraph-Cactus output was used to obtain subgenome aware genotype calls for the 71 isolates considered in the graph pangenome construction. Second, a mapping strategy was employed using the variation graph (vg) toolkit v1.54.0 (*45*) to generate genotypes in the population of 1,060 isolates (*27*). The graph pangenome was first converted to GBZ format using the *vg autoindex* command with the option *--workflow giraffe*. Snarls in the graph were identified using the *vg snarls* command. Paired-end Illumina reads were further mapped to the graph using *vg giraffe* with options *--fragment-mean 350 --fragment-stdev 100 -p - b fast --rescue-algorithm gssw* to obtain a GAM file. Read support from the GAM file was obtained using the *vg pack* command, with option *-Q 5* to ignore mapping and base quality below 5. Finally, a VCF file was generated for each sample using the *vg call* command with options *--genotype-snarls --all-snarls --snarls Graph.snarls.pb -k S.pack --sample S --ref-sample UMY321*, where *Graph.snarls.pb* is the snarls detected in the previous step, *S.pack* is the read support obtained previously and *S* is the sample name. The single-sample VCF files were merged into a single, multi-sample file using *bcftools merge* v1.18 (*46*) with option *--merge id*. Genotypes with DP < 2 were set as missing using *bcftools +setGT -- -t q -n . -e "FMT/DP>=2"*.

The obtained multi-sample VCF files were processed to keep only structural variants (> 50 bp) following these steps: First, loci without alleles larger than 50 bp were filtered out. Next, small variant alleles (< 50 bp) were converted to the reference allele to avoid considering small variants that overlap structural variants. Multiallelic loci were further split into biallelic records using *bcftools norm --multi-overlaps 0 -m -any*. To properly handle heterozygosity in the next step, diploid calls were split into two haploid calls. Similar SVs were merged using truvari v5.2.0 (*47*) with the command *truvari collapse --gt all*. Haploid calls were then merged back into diploid calls, and missing genotypes were set to the reference allele using the command *bcftools +setGT -- -t . -n 0* to ensure a conservative approach. Finally, SVs were annotated as (i) insertion, when the reference allele length is equal to 1, (ii) deletion, when the alternative allele length is equal to 1, and (iii) complex, when the length of both alleles is > 1. Any processing step not performed using bcftools was processed with custom Python scripts.

### Subgenome occurence of SVs

For the SV calls obtained from the Minigraph-Cactus output VCF file, the subgenome occurrence of SVs was determined from their presence among primary and subgenome assemblies. For the SV calls obtained from the graph mapping strategy, the subgenome occurrence of SVs was determined using a competitive mapping method. For each isolate, paired-end Illumina reads were mapped to the concatenation of the primary genome and corresponding acquired genome linear references, using bwa-mem2 v2.2.1 (*48*) with default parameters. Reads mapping to the primary and acquired subgenome references were separated using samtools v1.18 (*46*) with the command *samtools view --region-file contigs.txt -O BAM*, where *contigs.txt* corresponding to either the primary or the acquired subgenome contigs. *samtools fastq -1 Reads_1.fastq.gz -2 Reads_2.fastq.gz -0 /dev/null -s /dev/null -n* was then used. The SV calling pipeline was then performed independently on the primary and the acquired subgenome reads, with the only exception being that the *--ploidy 1* option was used for the acquired subgenome with the *vg call* command.

### Detection of acquired-to-primary genome conversion

For each allopolyploid isolate, regions of acquired-to-primary genome conversion were identified as a drop in nucleotide diversity, based on a short-read mapping strategy. Paired-end Illumina reads were mapped to the primary reference genome (*30*) using bwa-mem2 v2.2.1 (*48*). *Samtools sort* v1.18 (*46*) was used to sort the BAM files and GATK *AddOrReplaceReadGroups* command v4.5.0.0 (*49*) was used to add read groups. Variant calling was performed using Freebayes v1.3.2-dirty (*50*) with option *--ploidy 3*. SNPs were further extracted using bcftools v1.18 (*46*) and filtered for DP ≥ 3. Next, the number of SNPs within 50 kb windows sliding 1 kb was computed with VCFtools v0.1.16 (*51*) option *--SNPdensity 1000*. Peak and valley values were obtained from the kernel density of SNP across all windows. The discriminating threshold was chosen as the valley before the highest peak. A threshold of 19.16 SNP per kb was used to distinguish regions with the presence of both subgenomes from regions with acquired-to-primary genome conversion.

### Detection of MITE families

Identification of MITE elements was performed on all genome assemblies using MITE-tracker (*52*) and vsearch v2.7.1 (*53*) with option *--id 0.5*. The representative sequence of each MITE family (file families_nr.fasta) was aligned on SV sequences (inserted sequence for insertions, deleted sequence for deletions and both alleles for complex SVs) using blast v2.12.0 (*54*) with command *blastn -task blastn -outfmt "6 qseqid sseqid pident length mismatch gapopen qstart qend sstart send evalue bitscore qcovhsp qcovs" -query MITEsequences.fasta -subject SVsequences.fasta*. MITE-related SVs were defined by a percentage of identical position (*pident*) higher than 90% and a query coverage per subject (*qcovs*) higher than 80%.

### Structural diversity calculation

We defined the structural diversity 𝜋_!"_ as the average number of structural differences per site between two DNA sequences, similarly to nucleotide diversity:

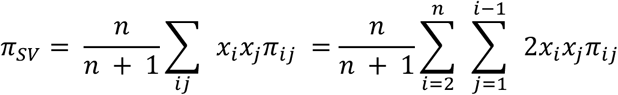

where 𝑥_$_ and 𝑥_%_ are the frequencies of the sequences 𝑖 and 𝑗, 𝜋_$%_ is the number of SVs between the two sequences and 𝑛 is the total number of sequences. The number of SVs was defined as the number of insertions, deletions and complex SVs overlapping the sequence of interest. Structural diversity was computed over the whole genome, for the complete population and per genetic cluster, and over 50 kb windows sliding 1 kb with a custom Python script.

### Genotype-phenotype local associations

The TPM for each gene was obtained from the Supplementary File 3 in ref. (*29*) and was used as a molecular phenotype. TPMs were normalized per gene using a rank-based inverse normal transformation manually implemented in R. A linear mixed-model regression was performed for each SV-gene pair separated by less than 50 kb, using the *lmer* function of the lmerTest R package v3.1-3 (*55*) with genetic cluster assignment as random effect. The effect size was computed as eta-squared using the *F_to_eta2* function of the effectsize R package v1.0.0 (*56*). *P* values were corrected using the FDR method implemented in the *adjust_pvalue* function of the rstatix R package v0.7.2 (*57*), and a threshold of 0.05 was used to detected significant associations.

## Code and data availability

Polished genome assemblies and reference graph pangenome, as well as code used to construct and analyze the graph are available in a Zenodo repository (*58*).

Temporary Zenodo link: https://zenodo.org/records/15740803?preview=1&token=eyJhbGciOiJIUzUxMiJ9.eyJpZCI6IjQ1ZDM3ZWVmLWY4YWUtNGFlMS05YzkxLTdjYmNiNDMxMjZiOSIsImRhdGEiOnt9LCJyYW5kb20iOiI1Y2YwZWY3M2ZmMWVjNTViOWRiY2Y2NDg5N2VhODkyYyJ9.BRGc-s7aDJE-mRKCrPEO8tKw7RuO4WqarAvoObhPgTqxDaBppQN9HjQedWFKztejMAVs3hYQxqRlDepqMKhuw

## Supplementary Material

**Figure S1.**
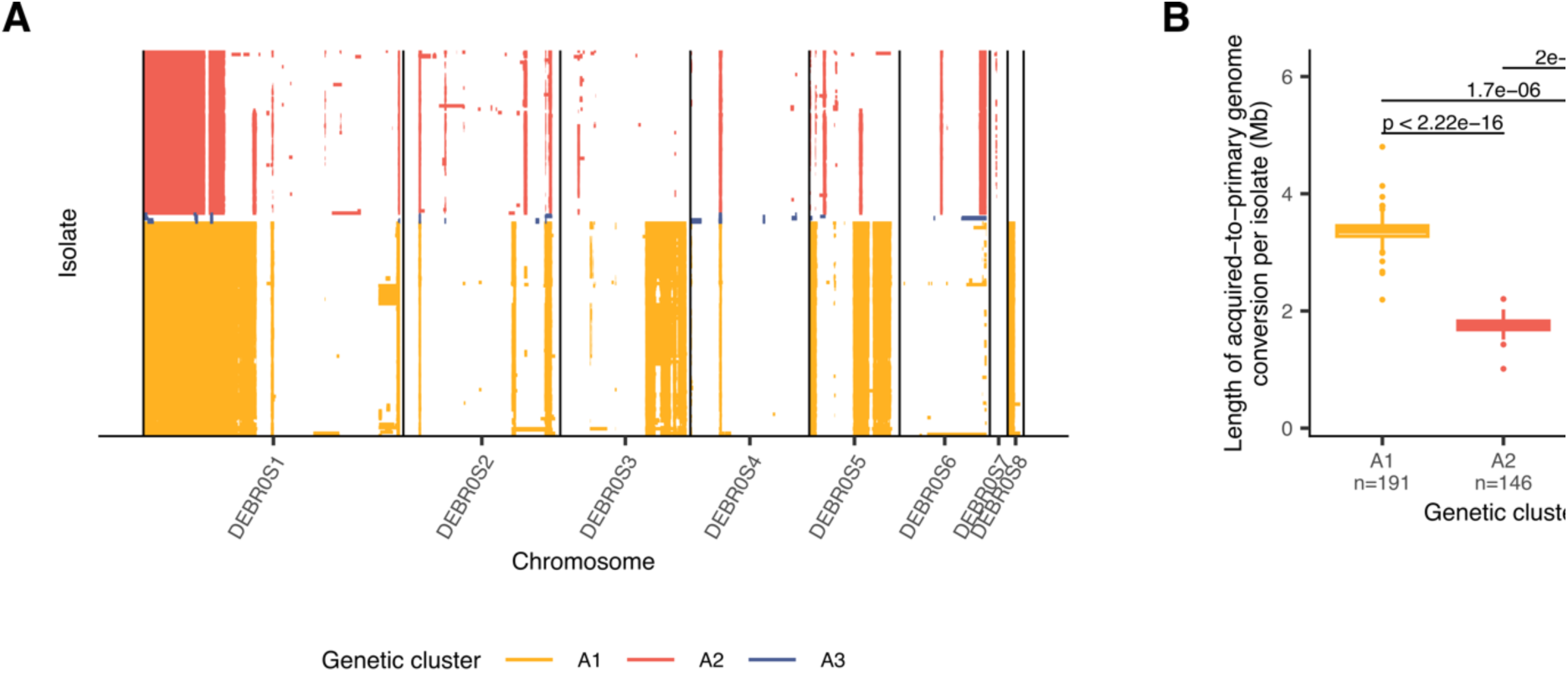
Acquired-to-primary subgenome conversion events. **A.** Regions of acquired-to-primary subgenome conversion per isolate. The genetic cluster is color-coded. **B.** Cumulative length regions with acquired-to-primary subgenome conversion per isolate. The middle bar of the box plots corresponds to the median; the upper and lower bounds correspond to the third and first quartiles, respectively. The whiskers correspond to the upper and lower bounds 1.5 times the interquartile range (IQR). The *P* values were computed using a two-sided rank-based Wilcoxon test without further adjustment.

**Figure S2.**
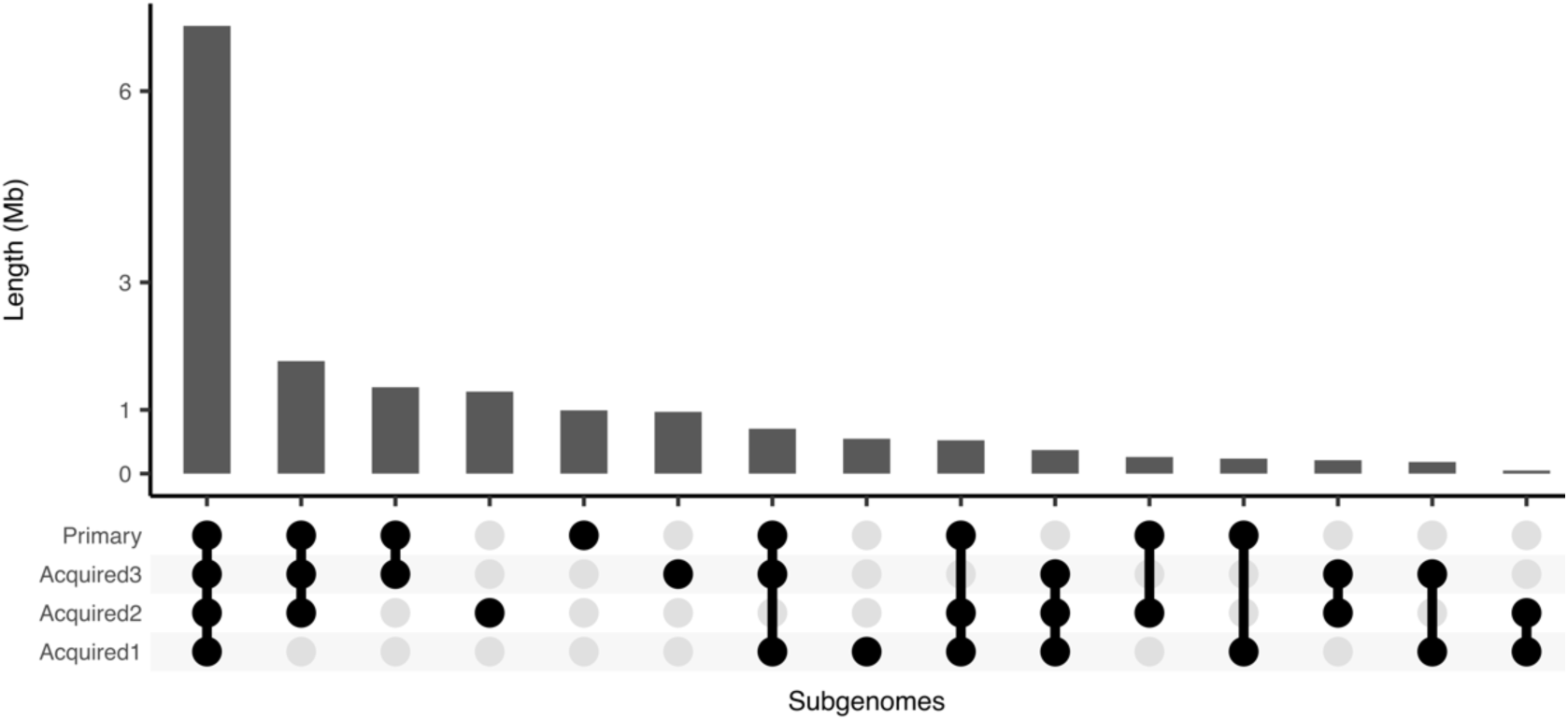
**Upset plot representing the cumulated length of shared segments across subgenome graphs.**

**Figure S3.**
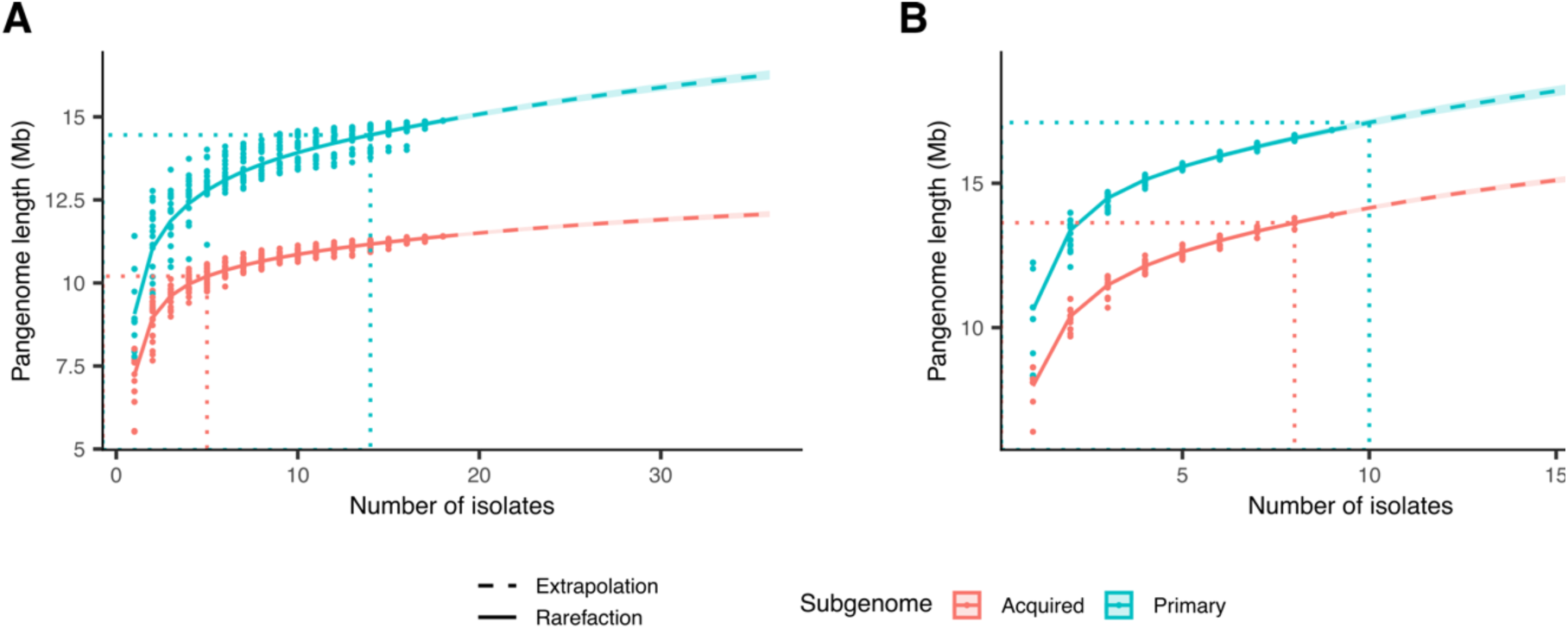
Subgenome rarefaction curves. **A.** Rarefaction curves for primary and acquired subgenomes among A1 isolates. **B.** Rarefaction curves for primary and acquired subgenomes in A2 isolates. The points represent observed rarefaction, the curves correspond to the non-parametric rarefaction model. The dotted lines indicate the pangenome length and number of isolates required to reach 80% of the pangenome saturation.

**Figure S4.**
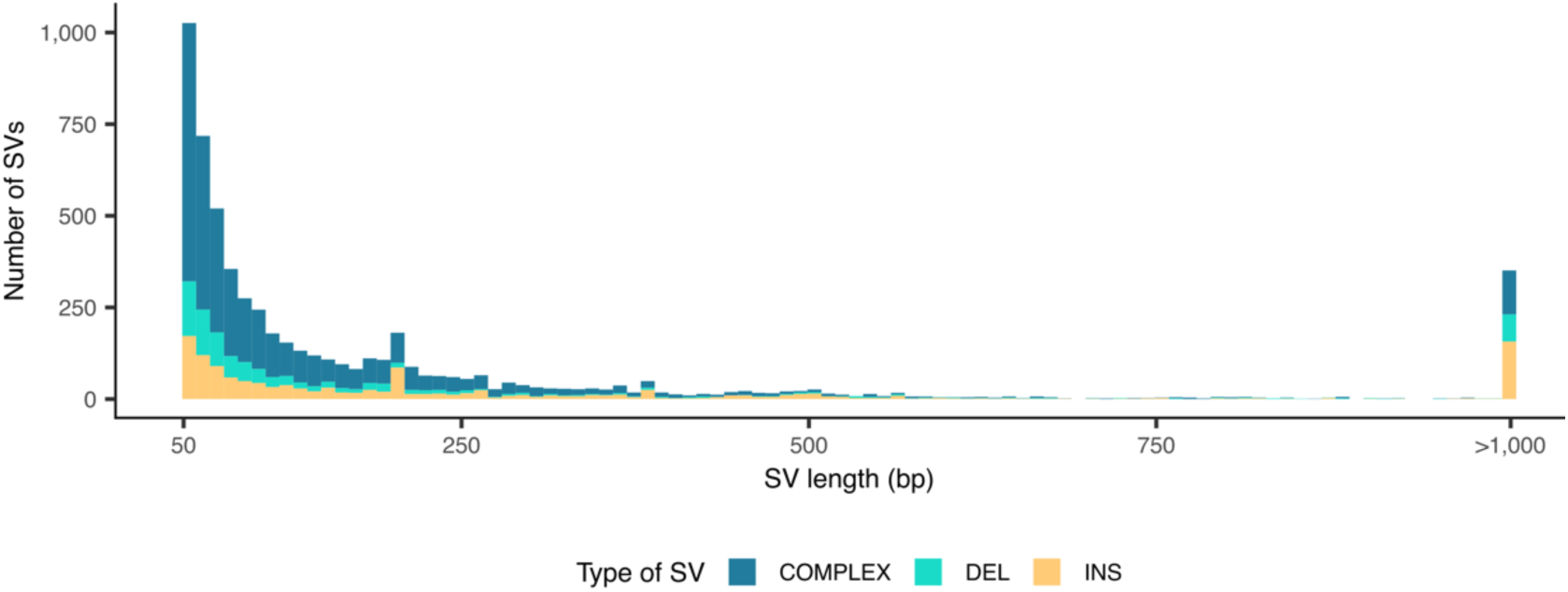
SV length distribution. The type of SV is color-coded.

**Figure S5.**
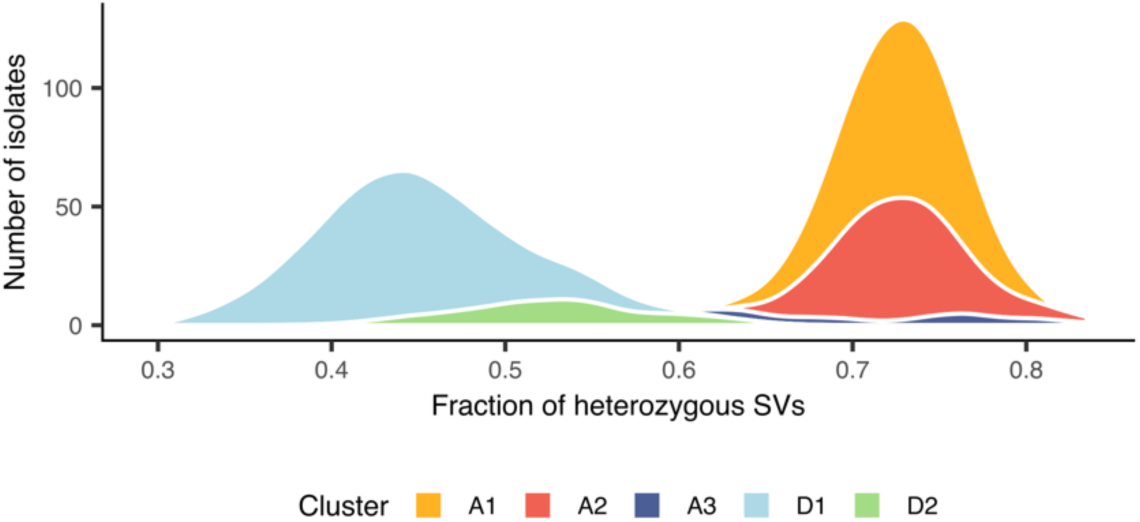
Distribution of the proportion of heterozygous SVs per isolate. Genetic clusters are color-coded.

