## Supplemental figures for "Graph-based pangenome analysis uncovers structural and functional impacts of allopolyploidization events"

**Supplementary Material**
**for**

### Graph-based pangenome analysis uncovers structural and functional impacts of allopolyploidization events

Victor Loegler[1], Anne Friedrich[1], Joseph Schacherer[1,2]

[1] Université de Strasbourg, CNRS, GMGM UMR 7156, Strasbourg, France
[2] Institut Universitaire de France (IUF), Paris, France

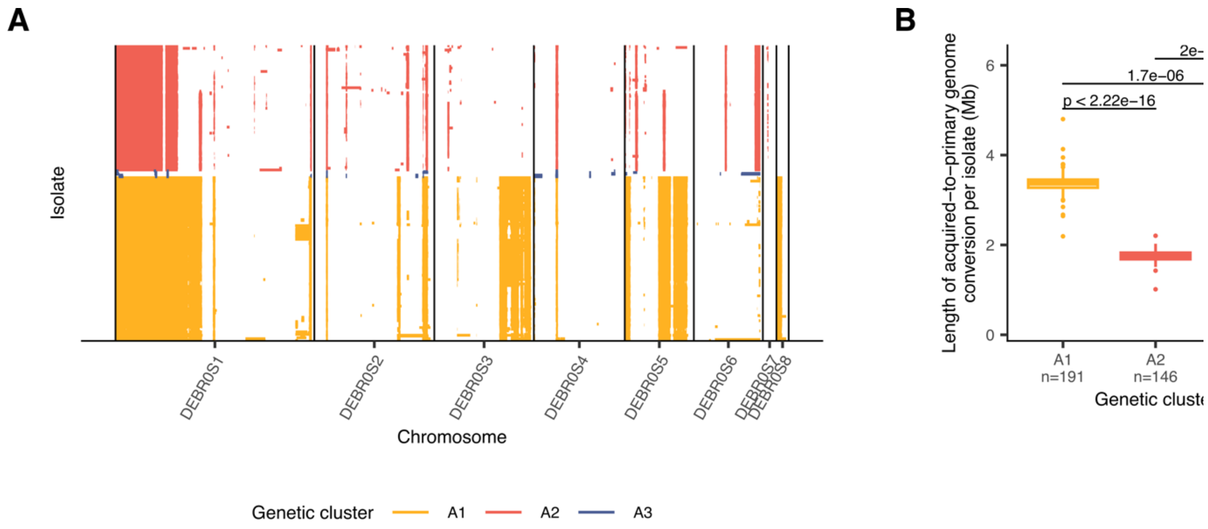


**Figure S1 | Acquired-to-primary subgenome conversion events. A.** Regions of acquired-to-primary subgenome conversion per isolate. The genetic cluster is color-coded. **B.** Cumulative length regions with acquired-to-primary subgenome conversion per isolate. The middle bar of the box plots corresponds to the median; the upper and lower bounds correspond to the third and first quartiles, respectively. The whiskers correspond to the upper and lower bounds 1.5 times the interquartile range (IQR). The *P* values were computed using a two-sided rank-based Wilcoxon test without further adjustment.

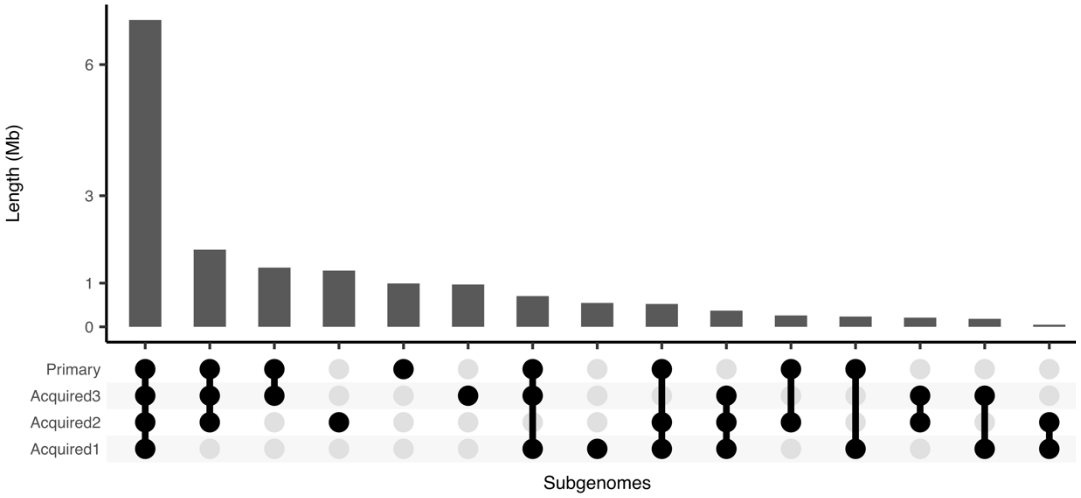


**Figure S2 | Upset plot representing the cumulated length of shared segments across subgenome graphs.**

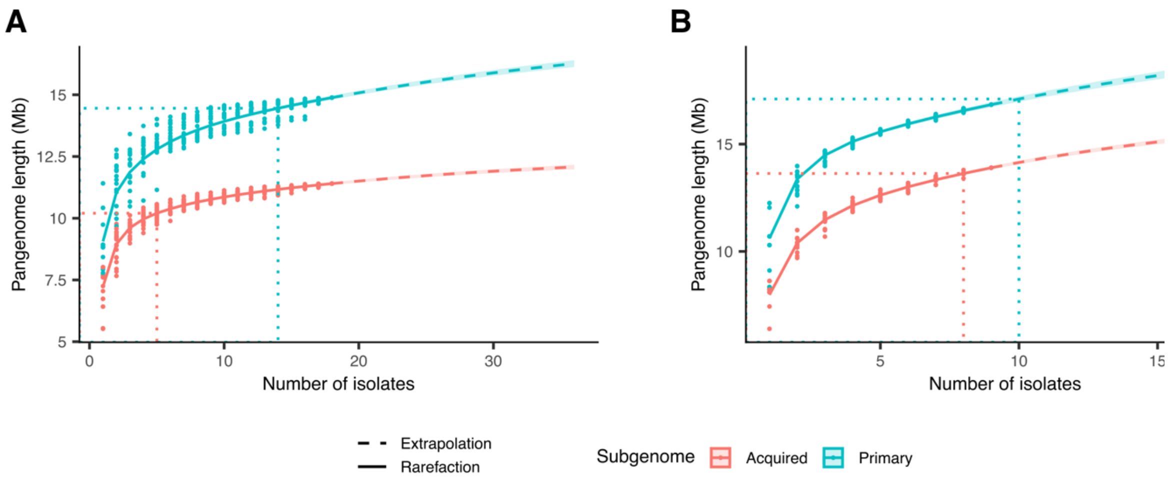


**Figure S3 | Subgenome rarefaction curves. A.** Rarefaction curves for primary and acquired subgenomes among A1 isolates. **B.** Rarefaction curves for primary and acquired subgenomes in A2 isolates. The points represent observed rarefaction, the curves correspond to the non-parametric rarefaction model. The dotted lines indicate the pangenome length and number of isolates required to reach 80% of the pangenome saturation.

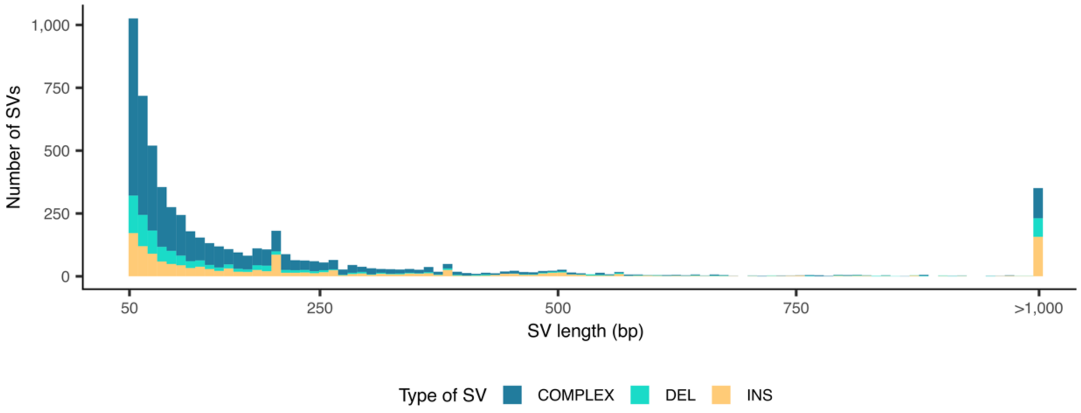


**Figure S4 | SV length distribution.** The type of SV is color-coded.

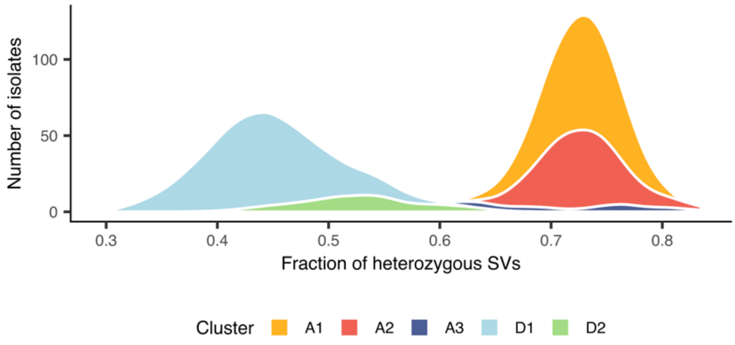


**Figure S5 | Distribution of the proportion of heterozygous SVs per isolate.** Genetic clusters are color-coded.
